# Lower adaptive immunity in invasive Egyptian geese compared to sympatric native waterfowls

**DOI:** 10.1101/2024.04.03.587943

**Authors:** Simone Messina, Hanna Prüter, Gábor Árpád Czirják, David Costantini

**Author notes:** **Corresponding author:** Simone Messina.

## Abstract

Successful invasive species increase their spreading success by trading-off nutritional and metabolic resources allocated to reproduction and range expansion with other costly body functions. One proposed mechanism for the reallocation of resources is a trade-off with the immune function and the regulation of oxidative status. Relying on a panel of blood-based markers of immune function and oxidative status quantified in an invasive species (Egyptian goose) and two native competing species (mallard and mute swan) in Germany, we tested the hypothesis that the invasive species would have (i) lower investment in immune function, (ii) lower levels of oxidative damage, and (iii) no higher antioxidant defences compared to the native species. We found lower levels of adaptive immune markers (lymphocytes and immunoglobulin Y), in the invasive species compared to the two native species. Reactive oxygen metabolites, a marker of oxidative damage, were also lower in Egyptian geese compared to the other species, while levels of antioxidants were generally similar. Mute swans showed the highest levels of heterophils, lysozymes and haemolysis among the three species, possibly due to its larger body mass. Our results point to a reduced investment in adaptive immune function in the invasive species as a possible resources-saving immunological strategy due to the loss of co-evolved parasites in the new colonised habitats, as observed in a previous study. A lower investment in immune function may benefit other energy-demanding activities, such as reproduction, dispersal, and territoriality.

**Research highlights:** Our results point to a reduced investment in adaptive immune function in invasive Egyptian geese compared to native and sympatric waterfowl species, as a possible resources-saving immunological strategy due to the loss of co-evolved parasites in the new colonised habitats.

## Introduction

The occurrence of biological invasions is increasing worldwide as a consequence of ongoing globalization. Successful invasive species engage in competition for vital resources with native species, sometimes even replacing native species in their natural habitat, and thus concurs with the ongoing biodiversity crisis (Mollot et al., 2017). Moreover, invaders can disrupt ecosystem function and facilitate the distribution of novel pathogens, including some with zoonotic potential, generating massive economic costs and challenges for public and animal health (Henry et al., 2023; Turbelin et al., 2023; Zhang et al., 2022).

Non-native introduced species need to establish and spread over the new region to become invasive. High population growth rate and superior dispersal ability are two main life-history traits that promote successful invasions (Sakai et al., 2001). However, increased need for nutritional and metabolic resources for particularly high reproductive investment and dispersion means that invaders need to trade-off those resources with other important body functions. Since invasive plants and animals may experience a loss of parasites during the invasion process (Dunn et al., 2012; Torchin et al., 2003), a reduced investment in the immune function might concur with the reallocation of resources (White & Perkins, 2012).

The immune system of vertebrates involves different pathways that require relatively higher or lower energetic costs. The evolution of increased competitive ability (EICA) hypothesis predicts that a loss of natural parasites in a habitat enables invasive species to colonise with reduced investment in defence functions and thus allocate more resources towards fitness-related traits (Blossey & Notzold, 1995). The EICA hypothesis was refined by Lee and Klasing (2004), who presented an analysis focused on the trade-offs between immune effectors in invasive vertebrates. The revised EICA hypothesis proposed that successful invasive species minimize systemic inflammatory responses that are triggered by strong innate or cell-mediated immune response to infectious agents, as they come at elevated energetic and immunopathological costs. Rather, a successful strategy for invaders to fight novel pathogens might be the increase of humoral defences, which includes both constitutive levels of natural antibodies and adaptive antibody responses (Lee & Klasing, 2004). Studies testing the EICA-hypothesis on vertebrates, however, show differential investment in immune functions between invasive and native species, and between high and low dispersers of an invasive species (revised in Cornet et al., 2016).

Maintenance of baseline immune function also requires investment of (micro)nutrients (Koutsos & Klasing, 2014). For example, an experimental study showed that nestling magpies (*Pica pica*) immunostimulated with methionine, a sulphur amino acid that enhances T-cell immune response, had reduced growth rates, but were capable to mount a stronger immune response to an immune challenge during the fledgling stage (Soler et al., 2003). In addition, Tschirren and Richner (2006) showed that nestling great tits (*Parus major*), whose immune system was boosted by extra provisioning of methionine, were able to grow at faster rate when exposed to hen fleas (*Ceratophyllus gallinae*), compared to control birds. These results indicate that the evolution of an optimal host defence is governed by a parasite-mediated allocation trade-off between fitness traits and immune function (Coon et al., 2014; White & Perkins, 2012). Accordingly, invasive species may take advantage of downregulating antibody-mediated responses (i.e. adaptive immunity) because they have lost their co-evolved parasites (Cornet et al., 2016).

Energy- and nutrient-based trade-off models provide valuable insight into the regulation of immune function in invaders. There are, however, other relevant physiological costs that can underlie immune investment. One important physiological mechanism linking immune response to individual fitness is the regulation of cellular oxidative status (Costantini & Møller, 2009; Hasselquist & Nilsson, 2012). Oxidative status is determined by levels of reactive oxygen species (ROS) generation, oxidative damage caused by ROS to biomolecules, and antioxidant molecules. During the phagocytosis of microbes, there is a rapid increase of ROS (respiratory burst) that enables phagocytic cells to degrade internalized particles and bacteria. Such an increase of ROS can induce oxidative stress if not sufficiently counterbalanced by antioxidants, with potential negative effects for the host fitness (Costantini, 2022).

Antioxidants prevent ROS from causing damages to biomolecules or even repair certain types of oxidative damage, and can also be upregulated to prevent disease or to protect the organism from immunopathology induced by the inflammatory response (Jena et al., 2023; Pham-Huy et al., 2008). Thus, upregulated baseline levels of endogenous antioxidants may help hosts to prevent a strong immune response and consequent oxidative stress. For example, white-browed sparrow weavers (*Plocepasser mahali*) with stronger superoxide dismutase (SOD) activity produced a smaller swelling response after an immune challenge (Cram et al., 2015). We are not aware of any studies that compared baseline levels of oxidative status markers between invasive and native species.

In this study, we compared the immune-physiological profile of the invasive Egyptian goose to that of two sympatric European waterfowl species. Specifically, we quantified a number of blood-based markers of immune function, oxidative status, and oxygen carrying capacity. The Egyptian goose is a successful invasive species which has been established in Europe since the 1980s and is currently expanding its range (Gyimesi & Lensink, 2012). Previous studies have found differences in immune markers and parasite prevalence between a native population of Egyptian geese (from Namibia) and the invasive population in our study area (Prüter et al., 2018, 2020). The other two waterfowl species included in this study are the mallard (*Anas platyrhynchos*) and the mute swan (*Cygnus olor*). They were chosen because resident and native to the study area, where they are sympatric with Egyptian geese, compete with them for territories, and are potentially exposed to the same pathogens. However, these three species also show differences (see Materials and Methods) that need to be considered in the interpretation of results.

Since we still have a poor understanding of how variation in immunological traits affects the success of biological invasions, making predictions on singular immune-physiological markers is challenging. We expect to find, however, downregulated immune effectors in the invasive species compared to the native species to save resources to be invested in other fitness traits. We also expect reduced oxidative damage in the invasive species due to its associated fitness costs. We do not expect higher levels of enzymatic antioxidants in the invasive species, compared to the native species, due to the energetic costs required for their synthesis.

## Materials and methods

### Study species and area

Mallard ducks and mute swans are waterfowl species native to Europe, while the Egyptian goose is native to sub-Saharan Africa and was introduced to Europe in the twentieth century. The invasion of the Egyptian goose started from the Netherlands, where they were introduced as ornamental species in urban parks. From the 1980s onwards, Egyptian geese also invade Germany where its population size increased rapidly. Nowadays, the Egyptian goose is the most common and widespread African waterfowl species in Europe (Gyimesi & Lensink, 2012). Mallard ducks, mute swans and Egyptian geese are resident, territorial and socially monogamous species. While mute swans and Egyptian geese show biparental care, mallards perform female-only parental care. Additionally, mallard populations show adult sex-ratios which are usually male-biased and forced extra-pair copulations frequently occur. Anseriformes are known for their aggressive behaviour towards conspecifics and other species, particularly during the breeding season, to protect their territory, nest and brood (Hohmann & Woog, 2021). All three species live in a variety of water habitats, with a clear preference for freshwater ponds, rivers and lakes (both natural and artificial) sometimes in close proximity to humans.

Adult individuals of the study species were live trapped in June and July 2016 in the Rhine and Mosel area in Western Germany (50.4° N, 7.6° E; Fig. S1). All individuals of this study were locally breeding and mostly in the phase of guiding goslings/ducklings at the time of sampling. Birds were caught using either loops or landing nets. Each bird was individually ringed, and body mass and sex were recorded. A blood sample from each bird was collected, either from the vena metatarsalia plantaris superficialis or the vena ulnaris, using a needle of 0.06 mm diameter. Blood samples were kept cool during field procedures, then centrifuged for 15 minutes to separate plasma from red blood cells which were stored separately in liquid nitrogen within 8 hours from blood collection. At the end of fieldwork, samples were stored at -80 °C until laboratory analyses.

Sample sizes for each assay were dependent on the total amount of plasma and red blood cells available from each individual, and therefore differ between the tests that were carried out (Table 1). Sampling was authorized in terms of animal welfare by the Landesuntersuchungsamt Rheinland-Pfalz (G 15-20-005) and Landesamt für Natur, Umwelt und Verbraucherschutz Nordrhein-Westfalen (LANUV) (84-08.04.2015.A266). The experimental procedures were approved by the Internal Committee for Ethics and Animal Welfare of the Leibniz Institute for Zoo and Wildlife Research (permit #2014-11-03).

**Table 1.** Sample size (n), mean and standard deviation (SD) for immunological effectors, and for haematocrit and markers of oxidative status.

| Immunological effectors | Egyptian goose |  | Mallard |  | Mute swan |  |
| --- | --- | --- | --- | --- | --- | --- |
|  | n | mean±SD | n | mean±SD | n | mean±SD |
| Lysozymes | 27 | 3.07±2 | 10 | 3.89±1.85 | 34 | 7.06±3.17 |
| Haemagglutination | 21 | 6.66±1.42 | 10 | 7.1±0.99 | 34 | 7.67±1.49 |
| Haemolysis | 21 | 5.76±1.13 | 10 | 6±0.94 | 34 | 6.94±1.59 |
| IgY | 23 | 0.55±0.19 | 10 | 0.81±0.22 | 36 | 0.86±0.2 |
| Bacteria Killing Ability | 27 | 83.61±24.77 | 10 | 84.01±25.77 | 27 | 47.61±23.04 |
| Basophils | 28 | 3.29±2.72 | 10 | 10.53±6.33 | 36 | 3.98±2.44 |
| Eosinophils | 28 | 26.99±13.97 | 10 | 33.14±19.78 | 36 | 25.21±15.48 |
| Heterophils | 28 | 6.98±4.26 | 10 | 11.17±11.27 | 36 | 16.69±9.83 |
| Lymphocytes | 28 | 35.18±14.61 | 10 | 51.38±19.6 | 36 | 49.87±18.99 |
| Monocytes | 28 | 15.1±6.2 | 10 | 15.57±10.01 | 36 | 10.89±7.24 |
| **Haematocrit and oxidative status** |  |  |  |  |  |  |
| Haematocrit | 24 | 44.79±8.5 | 7 | 40.57±8.81 | 21 | 39.9±4.51 |
| Non-enzymatic antioxidant capacity | 27 | 197.47±46.99 | 10 | 180.61±46.22 | 32 | 207.69±43.73 |
| Reactive Oxygen Metabolites | 23 | 0.33±0.15 | 8 | 0.68±0.26 | 32 | 0.44±0.22 |
| Protein Carbonyls | 18 | 43.23±17.97 | 8 | 40.27±14.61 | 30 | 35.31±18.05 |
| Glutathione peroxidase | 28 | 0.29±0.11 | 10 | 0.27±0.16 | 36 | 0.38±0.21 |
| Superoxide dismutase | 28 | 2.05±0.61 | 10 | 2.4±0.53 | 36 | 2.11±0.68 |

### Laboratory analyses

Several immunological assays were carried out to characterize both the cellular and humoral components of the adaptive and innate immune system of the study species (see supporting information). We used non-specific assays that had previously been applied to samples from a variety of free-living avian species, including different waterfowl (Bourgeon et al., 2010; Matson et al., 2006). Specifically, we counted basophils, eosinophils, heterophils and monocytes as effectors of cellular innate immunity. We quantified the amounts of lysozyme, bacteria killing ability against *E.coli* (BKA), haemagglutination and haemolysis as markers for humoral innate immunity. We measured the total immunoglobulin Y (IgY) concentration and the number of lymphocytes as markers of, respectively, humoral and cellular adaptive immunity (Table 1; Prüter et al., 2020).

In addition to the analyses of immune markers, we measured the haematocrit as a proxy for the oxygen carrying capacity of the blood (Ritchison, 2023). Further, we measured the total non-enzymatic antioxidant capacity (OXY-adsorbent assay), and the activity of antioxidant enzymes superoxide dismutase (SOD) and glutathione peroxidase (GPx), to estimate the antioxidant capacity of the blood. We measured the concentrations of protein carbonyls and reactive oxygen metabolites (d-ROMS test) to evaluate blood levels of oxidative damage. More details on the assays used for this study can be found in the Supplementary Material.

### Statistical analyses

The dataset included 74 actively reproducing individuals of which 28 were Egyptian geese, 10 were mallards, and 36 were mute swans. We collected blood samples from both female and male Egyptian geese (females = 12; males = 16) and mute swans (females = 21; males = 15), and only female mallards (Table 1).

We performed linear models (LMs) to assess the differences between species in each single marker. Therefore, for each model, we included one marker as a dependent variable and species as an independent variable. We always screened the model for the presence of outliers relying on the Cook’s distance. When we detected an outlier, this was removed and the model was run again. Then, we checked for normal distribution of residuals by means of Q-Q plots and Kolmogorov-Smirnov test (Bolker, 2008). If the distribution of model residuals was not normal, we log-transformed the dependent variables after which we performed again the model diagnostics (Fig. S2).

Subsequently, we implemented LMs testing for the interaction between species and sex on a subset of the original dataset including only Egyptian geese and mute swan (the mallard was excluded because we sampled only females). Following the previous procedures, we removed outliers when detected, we log-transformed the data when necessary; we performed the model diagnostics by visual inspection of Q-Q plots (Fig. S3) and Kolmogorov-Smirnov test. Significant interactions were explored by pairwise comparison through the function *emmeans* of the homonym package of R.

## Results

Results of models based on the full dataset (including outliers) are reported in Table S1. Here, we present results of models based on the dataset without outliers, because outliers can affect reliability and robustness of models (Knief & Forstmeier, 2021). Based on model diagnostic results, the distribution of residuals was normal for each model except that for Haemolysis (Kolmogorov-Smirnov test: statistic = 0.18, P-value = 0.03; Q-Q plots are shown in Fig. S2), even after the removal of outliers and log-trasformation of the dependent variable. Based on Knief and Forstmeier (2021), however, linear regression models are robust to violations of normality assumption and violating normality assumption is often unproblematic.

The outcomes of models show downregulated levels of lymphocytes and IgY (i.e. adaptive immunity) in Egyptian geese compared to both other species. Furthermore, Egyptian geese and mallards show lower levels of heterophils, lysozymes, and haemolysis, but higher levels of monocytes and BKA, than mute swans. In addition, Egyptian geese and mute swans show lower level of basophils than mallards (Table 2; Fig. 1).

**Fig. 1.**
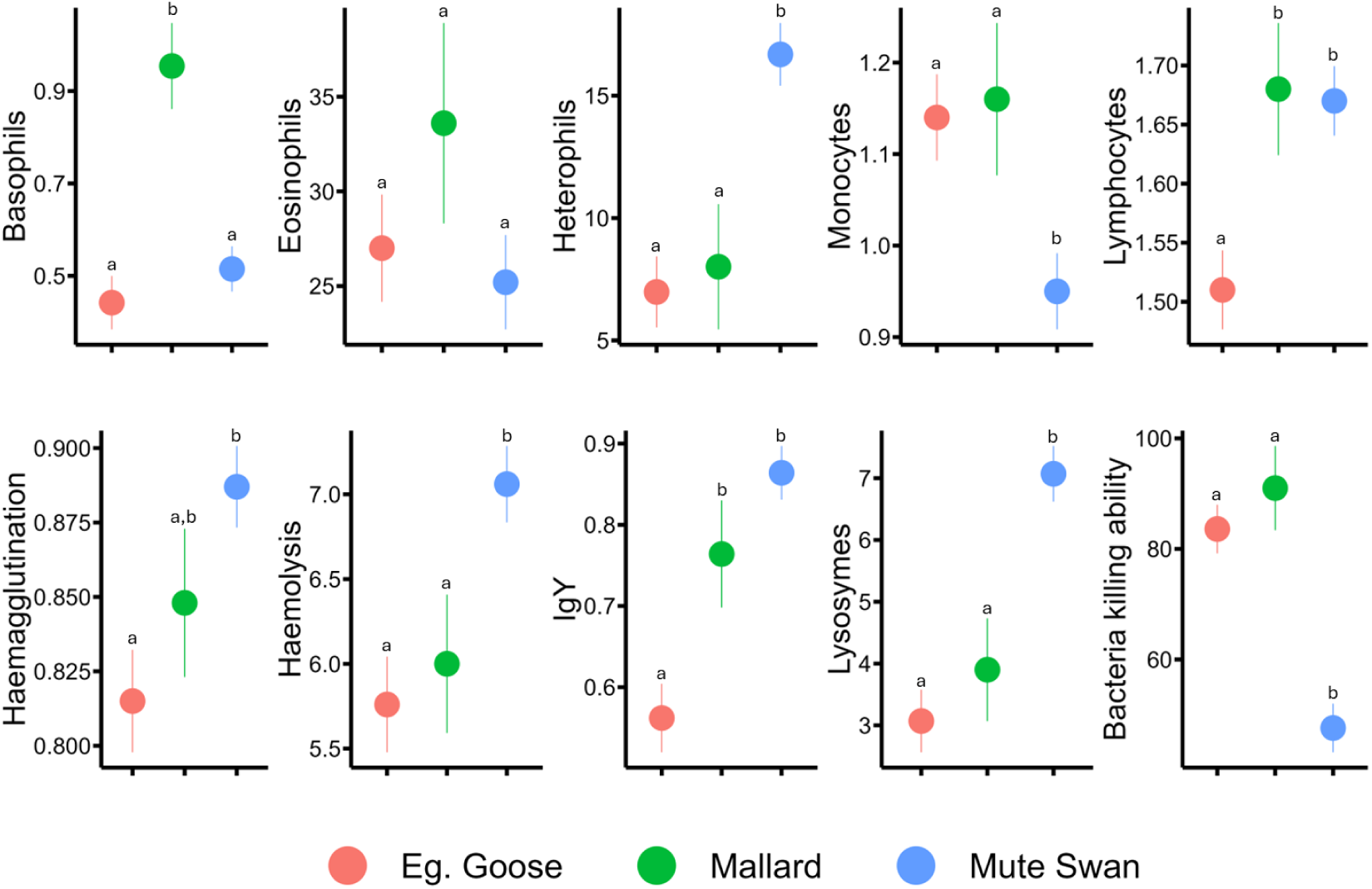
Estimated marginal means ± standard error of the mean immune markers level between species. Different superscripts (a and b) represent significant differences (*p* < 0.05).

**Table 2.** Results of LMs with outliers removed. We detected no outliers for lymphocytes, lysozymes, OXY and protein carbonyls, therefore results presented here for those markers are equal to those presented in Table S1. ALEG = Egyptian goose; ANPL = mallard; CYOL = mute swan. Significant P-values are indicated in bold.

| Marker | Variable tranformation | Contrast | Estimate | SE | t value | P-value |
| --- | --- | --- | --- | --- | --- | --- |
| Basophils | Log | ALEG - ANPL | -0.51 | 0.11 | -4.68 | **<0.01** |
|  |  | ALEG - CYOL | -0.08 | 0.08 | -0.97 | 0.34 |
|  |  | CYOL - ANPL | -0.44 | 0.11 | -4.17 | **<0.01** |
| Eosinophils | None | ALEG - ANPL | -6.59 | 5.99 | -1.10 | 0.28 |
|  |  | ALEG - CYOL | 1.79 | 3.77 | 0.47 | 0.64 |
|  |  | CYOL - ANPL | -8.38 | 5.85 | -1.43 | 0.16 |
| Heterophils | None | ALEG - ANPL | -1.03 | 2.94 | -0.35 | 0.73 |
|  |  | ALEG - CYOL | -9.71 | 1.93 | -5.03 | **<0.01** |
|  |  | CYOL - ANPL | 8.69 | 2.86 | 3.04 | **<0.01** |
| Lymphocytes | Log | ALEG - ANPL | -0.18 | 0.07 | -2.74 | **0.01** |
|  |  | ALEG - CYOL | -0.16 | 0.04 | -3.66 | **<0.01** |
|  |  | CYOL - ANPL | -0.02 | 0.06 | -0.24 | 0.81 |
| Monocytes | Log | ALEG - ANPL | -0.02 | 0.10 | -0.21 | 0.83 |
|  |  | ALEG - CYOL | 0.19 | 0.06 | 3.04 | **<0.01** |
|  |  | CYOL - ANPL | -0.21 | 0.09 | -2.27 | **0.03** |
| Haematocrit | None | ALEG - ANPL | 3.52 | 2.12 | 1.66 | 0.10 |
|  |  | ALEG - CYOL | 7.29 | 1.41 | 5.15 | **<0.01** |
|  |  | CYOL - ANPL | -3.76 | 2.12 | -1.77 | 0.08 |
| Lysozymes | None | ALEG - ANPL | -0.82 | 0.97 | -0.85 | 0.40 |
|  |  | ALEG - CYOL | -4.00 | 0.68 | -5.90 | **<0.01** |
|  |  | CYOL - ANPL | 3.17 | 0.94 | 3.36 | **<0.01** |
| Haemagglutination | Log | ALEG - ANPL | -0.03 | 0.03 | -1.09 | 0.28 |
|  |  | ALEG - CYOL | -0.07 | 0.02 | -3.30 | **<0.01** |
|  |  | CYOL - ANPL | 0.04 | 0.03 | 1.40 | 0.17 |
| Haemolysis | None | ALEG - ANPL | -0.24 | 0.50 | -0.48 | 0.63 |
|  |  | ALEG - CYOL | -1.30 | 0.36 | -3.60 | **<0.01** |
|  |  | CYOL - ANPL | 1.06 | 0.47 | 2.28 | **0.03** |
| IgY | None | ALEG - ANPL | -0.20 | 0.08 | -2.58 | **0.01** |
|  |  | ALEG - CYOL | -0.30 | 0.05 | -5.64 | **<0.01** |
|  |  | CYOL - ANPL | 0.10 | 0.07 | 1.36 | 0.18 |
| Bacteria killing ability | None | ALEG - ANPL | -7.36 | 8.80 | -0.84 | 0.41 |
|  |  | ALEG - CYOL | 36.00 | 6.22 | 5.78 | **<0.01** |
|  |  | CYOL - ANPL | -43.36 | 8.80 | -4.93 | **<0.01** |
| Non-enzymatic antioxidant capacity | None | ALEG - ANPL | 16.86 | 16.80 | 1.00 | 0.32 |
|  |  | ALEG - CYOL | -10.22 | 11.86 | -0.86 | 0.39 |
|  |  | CYOL - ANPL | 27.08 | 16.44 | 1.65 | 0.10 |
| Glutathione peroxidase | Log | ALEG - ANPL | 0.04 | 0.09 | 0.48 | 0.63 |
|  |  | ALEG - CYOL | -0.07 | 0.06 | -1.22 | 0.23 |
|  |  | CYOL - ANPL | 0.12 | 0.09 | 1.28 | 0.21 |
| Superoxide dismutase | None | ALEG - ANPL | -0.43 | 0.22 | -1.93 | 0.06 |
|  |  | ALEG - CYOL | -0.13 | 0.15 | -0.88 | 0.38 |
|  |  | CYOL - ANPL | -0.29 | 0.21 | -1.37 | 0.17 |
| Reactive Oxygen Metabolites | None | ALEG - ANPL | -0.34 | 0.08 | -4.18 | **<0.01** |
|  |  | ALEG - CYOL | -0.08 | 0.05 | -1.74 | 0.09 |
|  |  | CYOL - ANPL | -0.25 | 0.08 | -3.22 | **<0.01** |
| Protein carbonyls | None | ALEG - ANPL | 0.02 | 0.48 | 0.03 | 0.97 |
|  |  | ALEG - CYOL | 0.55 | 0.34 | 1.64 | 0.11 |
|  |  | CYOL - ANPL | -0.54 | 0.45 | -1.19 | 0.24 |

Regarding markers of oxidative status and haematocrit, mallards show higher levels of reactive oxygen metabolites compared to Egyptian geese and mute swan, and Egyptian geese show higher haematocrit compared to mute swan. We did not detect any differences among species for protein carbonyls and antioxidant markers (Table 2; Fig. 2).

**Fig. 2.**
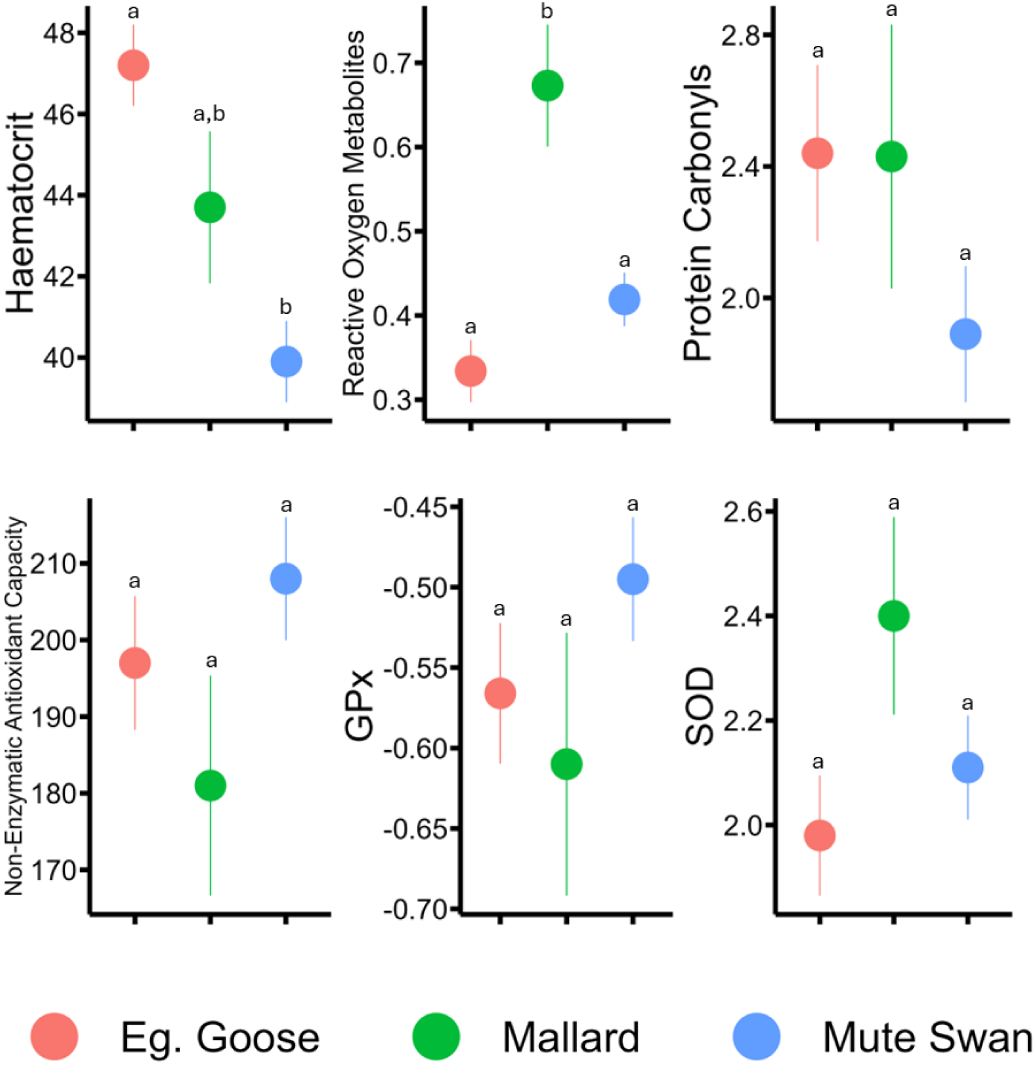
Estimated marginal means ± standard error of the mean immune markers level between species. GPx = glutathione peroxidase; SOD = superoxide dismutase. Different superscripts (a and b) represent significant differences (*p* < 0.05).

Finally, models testing for the interaction between species and sex showed a significant interaction for lysozymes only (Table S2). Pairwise comparison revealed higher levels of lysozymes in mute swans compared to Egyptian geese for both sexes, but a significant difference was on only detected for females (Table S3).

## Discussion

In this study we compared the baseline immune-physiological profile of an invasive species, the Egyptian goose, to that of two sympatric European waterfowl species, the mallard and the mute swan. We found lower levels of adaptive immune markers, i.e. lymphocytes and IgY, in the blood of the invasive species compared to the two native species. Our results also show a general similarity in the innate immune profile between Egyptian geese and mallards, with the only exception of basophils which are significantly lower in the invasive species. By contrast, mute swans show higher levels of heterophils, lysozymes and haemolysis compared to both Egyptian geese and mallards, and also the lowest levels of monocytes and BKA. Mallards also show the highest levels of reactive oxygen metabolites among the species (lowest levels were found in Egyptian geese, but the result was statistically significant only in respect to mallards).

### Adaptive immune effectors

Lymphocytes are one of the most abundant leucocytes in birds (Davis et al., 2008; Ritchison, 2023) and play important roles in both cell- and antibody-mediated adaptive immunity. Specifically, T-lymphocytes attack infected or abnormal cells inducing inflammation, and B-lymphocytes produce immunoglobulins (Guriec et al., 2018). Immunoglobulin Y (IgY) is the most abundant circulating immunoglobulin in birds and are released by B-lymphocytes (Ritchison, 2023). Thus, the lower levels of constitutive IgY in Egyptian geese are in agreement with the reduced number of lymphocytes, which point to a downregulated antibody-mediated immunity. According to Cornet et al. (2016), invasive hosts that are no more in contact with co-evolved (specialized) parasites should take advantage of downregulating antibody-mediated adaptive responses. In accordance, an experimental study on house finches (*Haemorhous mexicanus*) found higher levels of IgY in populations that co-evolved with the inoculated virus (*Mycoplasma gallisepticum*) compared to unexposed populations (Gates et al., 2021).

In accordance with our findings, recent studies found that German populations of Egyptian geese show lower prevalence of infections for a variety of blood parasites and macro-parasites compared to conspecifics from their native habitats and show lower seroprevalences of some common waterfowl viruses (Prüter et al., 2018, 2020). Such findings support the hypothesis that downregulated adaptive immunity in Egyptian geese in their invasive range may be due to reduced prevalences of infectious agents, which would be compatible with the predictions of the evolution of increased competitive ability hypothesis. On the other hand, a reduced investment in the less expensive antibody-mediated immunity is opposite to the predictions of the revised EICA-hypothesis. Further experimental studies are necessary to test the immune response of invasive Egyptian geese compared to that of native species.

### Innate immune effectors

We found that innate immunity effectors were generally similar between Egyptian geese and mallards. Differently, mute swan had significantly higher levels of heterophils, lysozymes and haemolysis, and lower levels of monocytes and BKA compared to the other species. These differences might be partly due to intrinsic characteristics of the species. For example, a recent multi-species study showed that avian heterophils increase with the body mass (Ruhs et al., 2020), and this might explain why we found higher levels of heterophils in mute swans compared to the other species. On the other hand, large species, such as mute swan, might be more exposed to parasites, and this can justify a different immune regulation (Downs et al., 2019).

Heterophils are the major phagocytic leucocytes and important mediators of acute inflammatory responses in birds (Scanes et al., 2022). Their higher levels in mute swans might indicate high prevalence of infections than in Egyptian geese and mallards. This result would agree with higher levels of haemagglutination and haemolysis in the mute swans (Table 1; Fig. 1). Haemagglutination and haemolysis reflect responses to foreign antigens driven by natural antibodies and the complement system, respectively (Matson et al., 2006). Natural antibodies are produced in absence of antigenic stimulation, independently from previous infections, whereas the complement system can be activated directly by pathogens or indirectly by immunoglobulins (Ochsenbein & Zinkernagel, 2000). Natural antibodies and the complement system are part of the innate immune system and act as a first line of defence against infections. Therefore, lower levels of haemagglutination and haemolysis might indicate that Egyptian geese and mallards have reduced prevalence of infections compared to mute swans. Mallards showed the highest level of basophils among the three species. Avian basophils, differently from those of mammals, are involved in inflammation reactions (Ritchison, 2023). The abundance of basophils in the blood of birds exhibits considerable variation among species, but Vinkler et al. (2010) found that basophil counts were higher in birds infected with blood parasites (Haemoproteus). Therefore, higher levels of basophils in mallards might be due to higher prevalence of haemosporidian infections other than different immune strategies compared to the other species. Interestingly, Egyptian goose is the only species showing no increase in any immunological mediator of inflammatory reactions. This observation supports the EICA-revised hypothesis, but further studies are needed to understand how baseline levels of circulating leukocytes might underlie different immunological strategies to fight pathogens.

Finally, our results show higher plasma BKA in Egyptian geese and mallards compared to mute swans. The mechanisms that allow plasma to kill bacteria are not well understood, but might be strongly dependent on antibacterial plasma proteins and peptides (Dugovich et al., 2019; Gao et al., 2015). Accordingly, Egyptian geese and mallards might invest relatively more energy in antimicrobial proteins and peptides, which come to lower (i) energetic costs compared to immune cells (e.g. heterophils) and (ii) fitness costs compared to inflammatory effectors.

### Oxidative status and haematocrit

We found higher levels of oxidative damage (reactive oxygen metabolites) in mallards compared to Egyptian geese and mute swans. In our study region, mallards are subject to increased competition and aggressive behaviour due to the presence of Egyptian geese (Hohmann & Woog, 2021), which might result in increased metabolic activity and ROS generation (Quque et al., 2022). It might also be that the higher pace of life of mallards compared to Egyptian geese and mute swans determines higher baseline generation of ROS. Egyptian geese exhibit the lowest levels of reactive oxygen metabolites (although only marginally significant as compared to mute swans). This result might indicate that invasive Egyptian geese are less prone to oxidative stress than native species. Reduced levels of oxidative stress may induce important benefits for the invasion success, such as higher reproductive success and longevity (Blount et al., 2016; Xia & Møller, 2018).

Finally, we found lower haematocrit in mute swans compared to the other species. Thus, variation in haematocrit does not seem to be relevant to achieve a successful invasion. Rather, our results are in agreement with previous studies that found lower haematocrit in larger bird species (Kostelecka-Myrcha et al., 1993).

### Conclusions

Our comparative analysis of multiple immune-physiological traits suggests that invasive Egyptian geese might strategically reduce their investment in adaptive immunity compared to sympatric related native species, because they are likely no longer exposed to the pathogens they encountered in their natural habitats. Thus, invasive species might invest less energy to sustain adaptive immune function, which might come at the benefit for other energy-demanding host traits, such as reproduction, dispersal, and territoriality. Besides energetic and nutritional costs, maintaining relatively higher innate immunity is beneficial during dispersal, or establishing in novel habitats, since invasive species mainly encounter novel pathogens. Future work is needed to understand how baseline differences in immunological parameters translate into trade-offs with fitness traits, such as reproduction or survival.

The results of our work also indicate species-specific differences in baseline immune status, supporting previous findings on the relationship between species’ body mass and heterophils. In addition, we also recognize that mallards show important behavioural, morphological, and pace-of-life differences with the other two species, which might contribute to among species variation in the regulation of oxidative status. On the other hand, physiological plasticity is an important driver of local acclimation and success of invasion (McCann et al., 2018). In accordance with Prüter et al. (2020), high levels of physiological plasticity in Egyptian geese might be one main mechanism underlying their successful invasion to Europe. Determining immune-physiological traits underlying the phenotype of successful invaders will foster our understanding of the proximate mechanisms through which species can successfully colonise novel environments and help to better predict novel alien species and potential mitigation strategies (Lennox et al., 2015). Future work should include further species that live simpatrically with the invasive Egyptian geese in order to better tease apart the effects of life-history variation from those induced by parasite exposure.

## Acknowledgements

S.M. was funded by the National Recovery and Resilience Plan (NRRP), Mission 4, Component 2, Investment 1.2 of the Italian Ministry for University and Research, funded by the European Union – NextGenerationEU, through the Young Researchers program (ID: SOE_0000189). The sample collection for this study was financially supported by the Ministry of Rhineland-Palatinate (Ministerium für Umwelt, Energie, Ernährung und Forsten, Project Nr: Gz. 105-63 313/2015-40). We thank Sönke Twietmeyer and Niklas Böhm for their great help with organizing and carrying out the field work. Also, we thank Lorena Derezanin, Sophie Ewert, Lea Jäger, Oliver Krone, Manuela Merling de Chapa, Katja Pohle, Felix Prüter, and Jannis Twietmeyer for their assistance during field or laboratory work. At last, we thank Rebecca Nagel for revising language and grammar of the manuscript. The laboratory analysis was supported by funds from the Leibniz Institute for Zoo and Wildlife Research, Berlin.

## Bibliography

Blossey, B., & Notzold, R. (1995). Evolution of increased competitive ability in invasive nonindigenous plants: A hypothesis. The Journal of Ecology, 83(5), 887. 10.2307/2261425

Blount, J. D., Vitikainen, E. I. K., Stott, I., & Cant, M. A. (2016). Oxidative shielding and the cost of reproduction. Biological Reviews, 91(2), 483–497. 10.1111/brv.12179

Bolker, B. M. (2008). Ecological models and data in R. Princeton university press.

Bourgeon, S., Kauffmann, M., Geiger, S., Raclot, T., & Robin, J.-P. (2010). Relationships between metabolic status, corticosterone secretion and maintenance of innate and adaptive humoral immunities in fasted re-fed mallards. Journal of Experimental Biology, 213(22), 3810–3818. 10.1242/jeb.045484

Coon, C. A. C., Brace, A. J., McWilliams, S. R., McCue, M. D., & Martin, L. B. (2014). Introduced and Native Congeners Use Different Resource Allocation Strategies to Maintain Performance during Infection. Physiological and Biochemical Zoology, 87(4), 559–567. 10.1086/676310

Cornet, S., Brouat, C., Diagne, C., & Charbonnel, N. (2016). Eco-immunology and bioinvasion: Revisiting the evolution of increased competitive ability hypotheses. Evolutionary Applications, 9(8), 952–962. 10.1111/eva.12406

Costantini, D. (2022). A meta-analysis of impacts of immune response and infection on oxidative status in vertebrates. Conservation Physiology, 10(1), coac018. 10.1093/conphys/coac018

Costantini, D., & Møller, A. P. (2009). Does immune response cause oxidative stress in birds? A meta-analysis. Comparative Biochemistry and Physiology Part A: Molecular & Integrative Physiology, 153(3), 339–344. 10.1016/j.cbpa.2009.03.010

Cram, D. L., Blount, J. D., York, J. E., & Young, A. J. (2015). Immune response in a wild bird is predicted by oxidative status, but does not cause oxidative stress. PLOS ONE, 10(3), e0122421. 10.1371/journal.pone.0122421

Davis, A. K., Maney, D. L., & Maerz, J. C. (2008). The use of leukocyte profiles to measure stress in vertebrates: A review for ecologists. Functional Ecology, 22(5), 760–772. 10.1111/j.1365-2435.2008.01467.x

Downs, C. J., Schoenle, L. A., Han, B. A., Harrison, J. F., & Martin, L. B. (2019). Scaling of Host Competence. Trends in Parasitology, 35(3), 182–192. 10.1016/j.pt.2018.12.002

Dugovich, B. S., Crane, L. L., Alcantar, B. B., Beechler, B. R., Dolan, B. P., & Jolles, A. E. (2019). Multiple innate antibacterial immune defense elements are correlated in diverse ungulate species. PLOS ONE, 14(11), e0225579. 10.1371/journal.pone.0225579

Dunn, A. M., Torchin, M. E., Hatcher, M. J., Kotanen, P. M., Blumenthal, D. M., Byers, J. E., Coon, C. A. C., Frankel, V. M., Holt, R. D., Hufbauer, R. A., Kanarek, A. R., Schierenbeck, K. A., Wolfe, L. M., & Perkins, S. E. (2012). Indirect effects of parasites in invasions. Functional Ecology, 26(6), 1262–1274. 10.1111/j.1365-2435.2012.02041.x

Gao, W., Xing, L., Qu, P., Tan, T., Yang, N., Li, D., Chen, H., & Feng, X. (2015). Identification of a novel cathelicidin antimicrobial peptide from ducks and determination of its functional activity and antibacterial mechanism. Scientific Reports, 5(1), 17260. 10.1038/srep17260

Gates, D. E., Staley, M., Tardy, L., Giraudeau, M., Hill, G. E., McGraw, K. J., & Bonneaud, C. (2021). Levels of pathogen virulence and host resistance both shape the antibody response to an emerging bacterial disease. Scientific Reports, 11(1), 8209. 10.1038/s41598-021-87464-9

Guriec, N., Bussy, F., Gouin, C., Mathiaud, O., Quero, B., Le Goff, M., & Collén, P. N. (2018). Ulvan activates chicken heterophils and monocytes through toll-like receptor 2 and toll-like receptor 4. Frontiers in Immunology, 9, 2725. 10.3389/fimmu.2018.02725

Gyimesi, A., & Lensink, R. (2012). Egyptian goose Alopochen aegyptiaca: An introduced species spreading in and from the Netherlands. Wildfowl, 62, 128–145.

Hasselquist, D., & Nilsson, J.-Å. (2012). Physiological mechanisms mediating costs of immune responses: What can we learn from studies of birds? Animal Behaviour, 83(6), 1303–1312. 10.1016/j.anbehav.2012.03.025

Henry, M., Leung, B., Cuthbert, R. N., Bodey, T. W., Ahmed, D. A., Angulo, E., Balzani, P., Briski, E., Courchamp, F., Hulme, P. E., Kouba, A., Kourantidou, M., Liu, C., Macêdo, R. L., Oficialdegui, F. J., Renault, D., Soto, I., Tarkan, A. S., Turbelin, A. J., … Haubrock, P. J. (2023). Unveiling the hidden economic toll of biological invasions in the European Union. Environmental Sciences Europe, 35(1), 43. 10.1186/s12302-023-00750-3

Hohmann, R., & Woog, F. (2021). How aggressive are Egyptian Geese Alopochen aegyptiaca? Interactions with Greylag Geese Anser anser and other birds in an urban environment. In Wildfowl. https://wildfowl.wwt.org.uk/index.php/wildfowl/article/view/2762

Jena, A. B., Samal, R. R., Bhol, N. K., & Duttaroy, A. K. (2023). Cellular Red-Ox system in health and disease: The latest update. Biomedicine & Pharmacotherapy, 162, 114606. 10.1016/j.biopha.2023.114606

Knief, U., & Forstmeier, W. (2021). Violating the normality assumption may be the lesser of two evils. Behavior Research Methods, 53(6), 2576–2590. 10.3758/s13428-021-01587-5

Kostelecka-Myrcha, A., Jaroszewicz, M., & Cholostiakow-Gromek, J. (1993). Relationship between the values of red blood indices and the body mass of birds. Acta Ornithologica, 28(1), 47–53.

Koutsos, E. A., & Klasing, K. C. (2014). Factors Modulating the Avian Immune System. In Avian Immunology (pp. 299–313). Elsevier. 10.1016/B978-0-12-396965-1.00017-0

Lee, K. A., & Klasing, K. C. (2004). A role for immunology in invasion biology. Trends in Ecology & Evolution, 19(10), 523–529. 10.1016/j.tree.2004.07.012

Lennox, R., Choi, K., Harrison, P. M., Paterson, J. E., Peat, T. B., Ward, T. D., & Cooke, S. J. (2015). Improving science-based invasive species management with physiological knowledge, concepts, and tools. Biological Invasions, 17(8), 2213–2227. 10.1007/s10530-015-0884-5

Matson, K. D., Cohen, A. A., Klasing, K. C., Ricklefs, R. E., & Scheuerlein, A. (2006). No simple answers for ecological immunology: Relationships among immune indices at the individual level break down at the species level in waterfowl. Proceedings of the Royal Society B: Biological Sciences, 273(1588), 815–822. 10.1098/rspb.2005.3376

McCann, S. M., Kosmala, G. K., Greenlees, M. J., & Shine, R. (2018). Physiological plasticity in a successful invader: Rapid acclimation to cold occurs only in cool-climate populations of cane toads (Rhinella marina). Conservation Physiology, 6(1). 10.1093/conphys/cox072

Mollot, G., Pantel, J. H., & Romanuk, T. N. (2017). The Effects of Invasive Species on the Decline in Species Richness. In Advances in Ecological Research (Vol. 56, pp. 61–83). Elsevier. 10.1016/bs.aecr.2016.10.002

Ochsenbein, A. F., & Zinkernagel, R. M. (2000). Natural antibodies and complement link innate and acquired immunity. Immunology Today, 21(12), 624–630. 10.1016/S0167-5699(00)01754-0

Pham-Huy, L. A., He, H., & Pham-Huy, C. (2008). Free radicals, antioxidants in disease and health. International Journal of Biomedical Science: IJBS, 4(2), 89–96.

Prüter, H., Czirják, G. Á., Twietmeyer, S., Harder, T., Grund, C., Mühldorfer, K., & Lüschow, D. (2018). Sane and sound: A serologic and molecular survey for selected infectious agents in neozootic Egyptian geese (Alopochen aegyptiacus) in Germany. European Journal of Wildlife Research, 64(6), 71. 10.1007/s10344-018-1231-9

Prüter, H., Franz, M., Twietmeyer, S., Böhm, N., Middendorff, G., Portas, R., Melzheimer, J., Kolberg, H., Von Samson-Himmelstjerna, G., Greenwood, A. D., Lüschow, D., Mühldorfer, K., & Czirják, G. Á. (2020). Increased immune marker variance in a population of invasive birds. Scientific Reports, 10(1), 21764. 10.1038/s41598-020-78427-7

Quque, M., Ferreira, C., Sosa, S., Schull, Q., Zahn, S., Criscuolo, F., Bleu, J., & Viblanc, V. A. (2022). Cascading Effects of Conspecific Aggression on Oxidative Status and Telomere Length in Zebra Finches. Physiological and Biochemical Zoology, 95(5), 416–429. 10.1086/721252

Ritchison, G. (2023). In a Class of Their Own: A Detailed Examination of Avian Forms and Functions. Springer International Publishing. 10.1007/978-3-031-14852-1

Ruhs, E. C., Martin, L. B., & Downs, C. J. (2020). The impacts of body mass on immune cell concentrations in birds. Proceedings of the Royal Society B: Biological Sciences, 287(1934), 20200655. 10.1098/rspb.2020.0655

Sakai, A. K., Allendorf, F. W., Holt, J. S., Lodge, D. M., Molofsky, J., With, K. A., Baughman, S., Cabin, R. J., Cohen, J. E., Ellstrand, N. C., McCauley, D. E., O’Neil, P., Parker, I. M., Thompson, J. N., & Weller, S. G. (2001). The population biology of invasive species. Annual Review of Ecology and Systematics, 32(1), 305–332. 10.1146/annurev.ecolsys.32.081501.114037

Scanes, C. G., Dridi, S., & Sturkie, P. D. (A c. Di). (2022). Sturkie’s avian physiology (Seventh edition). Academic Press, an imprint of Elsevier.

Soler, J. J., Neve, L. D., Pérez–Contreras, T., Soler, M., & Sorci, G. (2003). Trade-off between immunocompetence and growth in magpies: An experimental study. Proceedings of the Royal Society of London. Series B: Biological Sciences, 270(1512), 241–248. 10.1098/rspb.2002.2217

Torchin, M. E., Lafferty, K. D., Dobson, A. P., McKenzie, V. J., & Kuris, A. M. (2003). Introduced species and their missing parasites. Nature, 421(6923), 628–630. 10.1038/nature01346

Tschirren, B., & Richner, H. (2006). Parasites shape the optimal investment in immunity. Proceedings of the Royal Society B: Biological Sciences, 273(1595), 1773–1777. 10.1098/rspb.2006.3524

Turbelin, A. J., Cuthbert, R. N., Essl, F., Haubrock, P. J., Ricciardi, A., & Courchamp, F. (2023). Biological invasions are as costly as natural hazards. Perspectives in Ecology and Conservation, 21(2), 143–150. 10.1016/j.pecon.2023.03.002

Vinkler, M., Schnitzer, J., Munclinger, P., Votýpka, J., & Albrecht, T. (2010). Haematological health assessment in a passerine with extremely high proportion of basophils in peripheral blood. Journal of Ornithology, 151(4), 841–849. 10.1007/s10336-010-0521-0

White, T. A., & Perkins, S. E. (2012). The ecoimmunology of invasive species. Functional Ecology, 26(6), 1313–1323. 10.1111/1365-2435.12012

Xia, C., & Møller, A. P. (2018). Long-lived birds suffer less from oxidative stress. Avian Research, 9(1), 41. 10.1186/s40657-018-0133-6

Zhang, L., Rohr, J., Cui, R., Xin, Y., Han, L., Yang, X., Gu, S., Du, Y., Liang, J., Wang, X., Wu, Z., Hao, Q., & Liu, X. (2022). Biological invasions facilitate zoonotic disease emergences. Nature Communications, 13(1), 1762. 10.1038/s41467-022-29378-2

