## Supplementary material for "Lower adaptive immunity in invasive Egyptian geese compared to sympatric native waterfowls": Supporting_Material.docx

**Immune-physiological assays**

*Leucocytes count*. To count leucocytes, blood smears were prepared, air-dried and stained using Giemsa- and May-Grünwald staining. Smears were examined at 1,000 fold magnification with oil immersion and the relative number and types of leucocytes were assessed by counting 100 leucocytes. The number of white blood cells of different types was expressed per 10^4^ erythrocytes (Pap et al., 2015).

*Immunoglobulin Y.* Total IgY, the avian equivalent to mammalian IgG, was measured using a sensitive ELISA

with commercial anti-chicken antibodies (Bourgeon et al., 2010; Martínez et al., 2003). Ninety-six-well high-binding ELISA plates (82.1581.200, Sarstedt) were coated with 100 μl of diluted serum sample (2 samples per bird 1:16,000 diluted in carbonate–bicarbonate buffer) and incubated first for 1 h at 37 °C and then overnight at 4 °C. After incubation, the plates were washed with a 200 μl solution of phosphate buffer saline and PBS–Tween, before 100 μl of a solution of 1% gelatine in PBS–Tween was added. Plates were then incubated at 37 °C for 1 h, washed with PBS–Tween and 100 μl of polyclonal rabbit anti-chicken IgY conjugated with peroxidase (A-9046, Sigma) at 1:250 (v/v) was added. Following 2 h incubation at 37 °C, the plates were washed again with PBS–Tween three times. After washing, 100 μl of revealing solution [peroxide diluted 1:1000 in ABTS (2,20-azino-bis- (3-ethylbenzthiazoline-6-sulphonic acid))] was added, and the plates were incubated for 1 h at 37 °C. The final absorbance was measured at 405 nm using a photometric microplate reader (μQuant Microplate Spectrophotometer, Biotek) and subsequently defined as total serum IgY levels (Bourgeon & Raclot, 2006).

*Lysozyme.* To measure lysozyme concentration in serum, we used the lysoplate assay (Giraudeau et al., 2010): 25 μl serum were inoculated in the test holes of a 1% Noble agar gel (A5431, Sigma) containing 50 mg/100 ml lyophilized *Micrococcus lysodeikticus* (M3770, Sigma), a bacteria which is particularly sensitive to lysozyme concentration. Crystalline hen egg white lysozyme (L6876, Sigma) (concentration: 1, 1.25, 2.5, 5, 6.25, 10, 12.5, 20 and 25 μg/ml) was used to prepare a standard curve for each plate. Plates were incubated at room temperature (25–27 °C) for 20 h. During this period, as a result of bacterial lysis, a clear zone developed in the area of the gel surrounding the sample inoculation site. The diameters of the cleared zones are proportional to the log of the lysozyme concentration. This area was measured three times digitally using the software ImageJ (version 1.48, http://image j.nih.gov/ij/) and the mean was converted to a semi-logarithmic plot into hen egg lysozyme equivalents (HEL equivalents, expressed in μg/mL) according to the standard curve (Rowe et al., 2013).

*Haemolysis–haemagglutination assay.* The levels of the natural antibodies and complement were assessed by using a haemolysis–haemagglutination assay as described by (Matson et al., 2005) adjusted to the limited volume of serum. After pipetting 15 μl of serum into the first two columns of a U-shaped 96-well microtitre plate, 15 μl sterile PBS were added to columns 2–12. The content of the second column wells was serially diluted (1:2) until the 11th column, resulting in a dilution series for each sample from 1/1 to 1/1024. The last column of the plate was used as negative controls, containing PBS only. Fifteen μl of 1% rabbit red blood cells (supplied as 50% whole blood, 50% Alsever’s solution, Envigo) suspension was added to all wells and incubated at 37 °C for 90 min. After incubation, in order to increase the visualisation of agglutination, the plates were tilted at a 45° angle at room temperature. Agglutination and lysis, which reflect the activity of the natural antibodies and the interaction between these antibodies and complement (Matson et al., 2005; Pap et al., 2010), was recorded after 20 and 90 min, respectively. Haemagglutination is characterised by the appearance of clumped red blood cells, as a result of antibodies binding multiple antigens, while during haemolysis, the red blood cells are destroyed. Titres of the natural antibodies and complement were given as the log2 of the reciprocal of the highest dilution of serum showing positive haemagglutination or lysis, respectively (Matson et al., 2005; Pap et al., 2015).

*Bacteria killing ability*. We measured the soluble aspects of the constitutive innate immunity by assessing the *in vitro* bacterial killing activity of the serum against *Escherichia coli* following the original version of the method (Tieleman et al., 2005). Plasma samples were diluted 1:10 in PBS (pH=7.4) and to each diluted sample (140 μl) we added 10 μl of a suspension of live *E. coli* (ATCC #8739). The bacterial suspension was adjusted to a concentration of ~200 colonies per 50 μl of diluted plasma-bacteria mixture. After incubation, for 30 min at 40ºC (avian body temperature), 50 μl of the plasma-bacteria mixture was spread aliquots onto Tryptic Soy Agar plates (#CP70.1, Carl Roth GmbH) in duplicate, and the plates were incubated overnight at 37ºC. To obtain the initial number of bacteria that we had before starting to interact with the plasma, we diluted 140 μl media alone with bacterial suspension and plated immediately. On the following day the colony-forming units were counted and the bacterial killing activity was defined as the percent of the killed bacteria, which was calculated as 1 – (average of the viable bacteria after incubation / the initial number of bacteria). The average was calculated from two plates per sample.

*Haematocrit.*

We collected whole blood in heparinized microhaematocrit capillary tubes and centrifuged the samples directely after collection, thus separating plasma from cellular blood. Hct levels were measured for each full capillary as the percentage of packed red blood cells over the total blood sample.

*Protein carbonyls.* Protein carbonyls (marker of oxidative protein damage) were measured using the Protein Carbonyl Colorimetric assay (Cayman Chemical Company, Ann Arbor, USA). This assay is based on the colorimetric method proposed by Levine et al. (1990). A same volume of plasma was used for all samples and the amount of carbonyls was standardised by the plasma protein concentration according to manufacturer's instructions. Protein carbonyls were derivatized to 2,4-dinitrophenylhydrazone by reaction with 2,4-dinitrophenylhydrazine (DNPH). The absorbance was read at 370 nm. The extinction coefficient for DNPH (0.022/μM/cm) was used to calculate the concentration of protein carbonyls, which was expressed as nmol/mg protein (amount of carbonyls generated per unit of protein) or as total nmol/ml obtained by multiplying the concentration of carbonyls by the concentration of plasma proteins (i.e., total amount of carbonyls in the sample, which is also dependent on the amount of substrates available, i.e., proteins). The metric is expressed as total amount of carbonyls that occurs in the tissue (ml), which is influenced by the amount of proteins available. This metric is important because accumulation of carbonyls is detrimental for the cells (Halliwell & Gutteridge, 2015).

*Reactive oxygen metabolites*. Serum reactive oxygen metabolites, a marker of intermediate oxidative damage generated early in the oxidative cascade, were measured in duplicate using the d-ROMs assay (Diacron International, Grosseto, Italy) and values were expressed as mM H_2_O_2_ equivalents.

*Non-enzymatic antioxidant capacity.* The OXY-Adsorbent test (Diacron International, Italy) was used to quantify the non-enzymatic antioxidant capacity of plasma against HOCl. Values were expressed as mM of HOCl neutralised.

*Glutathione peroxidase and superoxide dismutase*. The Ransel assay (RANDOX Laboratories, UK) was used to measure the activity of the antioxidant enzyme glutathione peroxidase (GPx) in haemolysates (red blood cells diluted with distilled water). Values were expressed as Units of GPx per mg of proteins. The activity in red blood cells of the enzyme superoxide dismutase (SOD), which prevents oxidation due to superoxide radical, was measured in duplicate using the Ransod assay (RANDOX Laboratories, Crumlin, UK) and was expressed as units of SOD per mg of proteins. The Bradford protein assay (Bio-Rad Laboratories, Hercules, USA) with bovine albumin as a reference standard was used to measure the concentration of proteins in both plasma samples and haemolysates.

**Bibliography**

Bourgeon, S., Kauffmann, M., Geiger, S., Raclot, T., & Robin, J.-P. (2010). Relationships between metabolic status, corticosterone secretion and maintenance of innate and adaptive humoral immunities in fasted re-fed mallards. *Journal of Experimental Biology*, *213*(22), 3810–3818. https://doi.org/10.1242/jeb.045484

Bourgeon, S., & Raclot, T. (2006). Corticosterone selectively decreases humoral immunity in female eiders during incubation. *Journal of Experimental Biology*, *209*(24), 4957–4965. https://doi.org/10.1242/jeb.02610

Giraudeau, M., Czirják, G. Á., Duval, C., Bretagnolle, V., Eraud, C., McGraw, K. J., & Heeb, P. (2010). Effect of Restricted Preen-Gland Access on Maternal Self Maintenance and Reproductive Investment in Mallards. *PLoS ONE*, *5*(10), e13555. https://doi.org/10.1371/journal.pone.0013555

Halliwell, B., & Gutteridge, J. M. C. (2015). *Free radicals in biology and medicine* (Fifth edition). Oxford University Press.

Martínez, J., Tomás, G., Merino, S., Arriero, E., & Moreno, J. (2003). Detection of serum immunoglobulins in wild birds by direct ELISA: A methodological study to validate the technique in different species using antichicken antibodies. *Functional Ecology*, *17*(5), 700–706. https://doi.org/10.1046/j.1365-2435.2003.00771.x

Matson, K. D., Ricklefs, R. E., & Klasing, K. C. (2005). A hemolysis–hemagglutination assay for characterizing constitutive innate humoral immunity in wild and domestic birds. *Developmental & Comparative Immunology*, *29*(3), 275–286. https://doi.org/10.1016/j.dci.2004.07.006

Pap, P. L., Czirják, G. Á., Vágási, C. I., Barta, Z., & Hasselquist, D. (2010). Sexual dimorphism in immune function changes during the annual cycle in house sparrows. *Naturwissenschaften*, *97*(10), 891–901. https://doi.org/10.1007/s00114-010-0706-7

Pap, P. L., Vágási, C. I., Vincze, O., Osváth, G., Veres-Szászka, J., & Czirják, G. Á. (2015). Physiological pace of life: The link between constitutive immunity, developmental period, and metabolic rate in European birds. *Oecologia*, *177*(1), 147–158. https://doi.org/10.1007/s00442-014-3108-2

Rowe, M., Czirják, G. Á., Lifjeld, J. T., & Giraudeau, M. (2013). Lysozyme-associated bactericidal activity in the ejaculate of a wild passerine: Lysozyme in the Ejaculate of a Wild Bird. *Biological Journal of the Linnean Society*, *109*(1), 92–100. https://doi.org/10.1111/bij.12044

Tieleman, B. I., Williams, B. J., Ricklefs, E. R., & Klasing, C. K. (2005). Constitutive innate immunity is a component of the pace-of-life syndrome in tropical birds. *Proceedings of the Royal Society B: Biological Sciences*, *272*(1573), 1715–1720. https://doi.org/10.1098/rspb.2005.3155

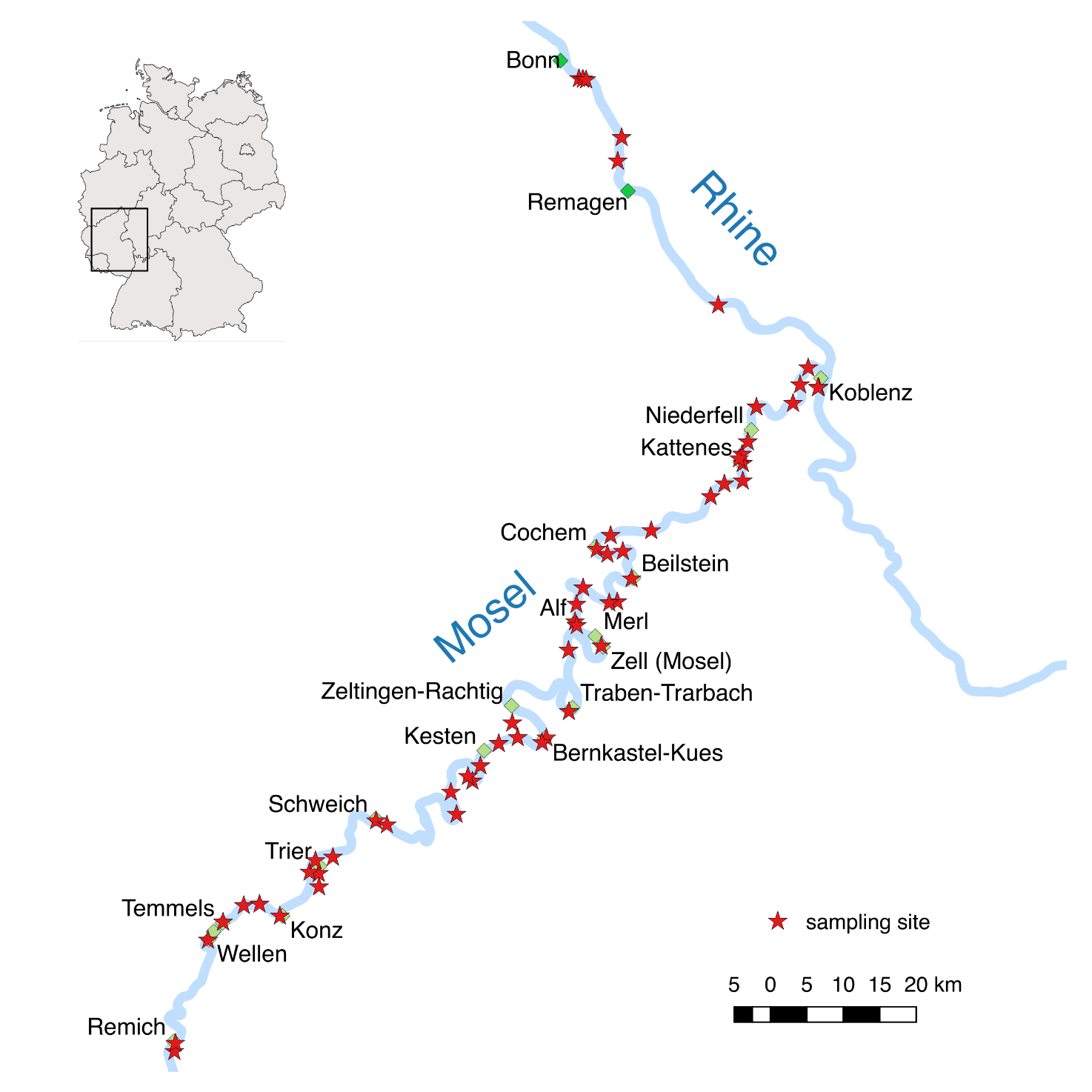

Fig. S1 – Study area and sampling sites.

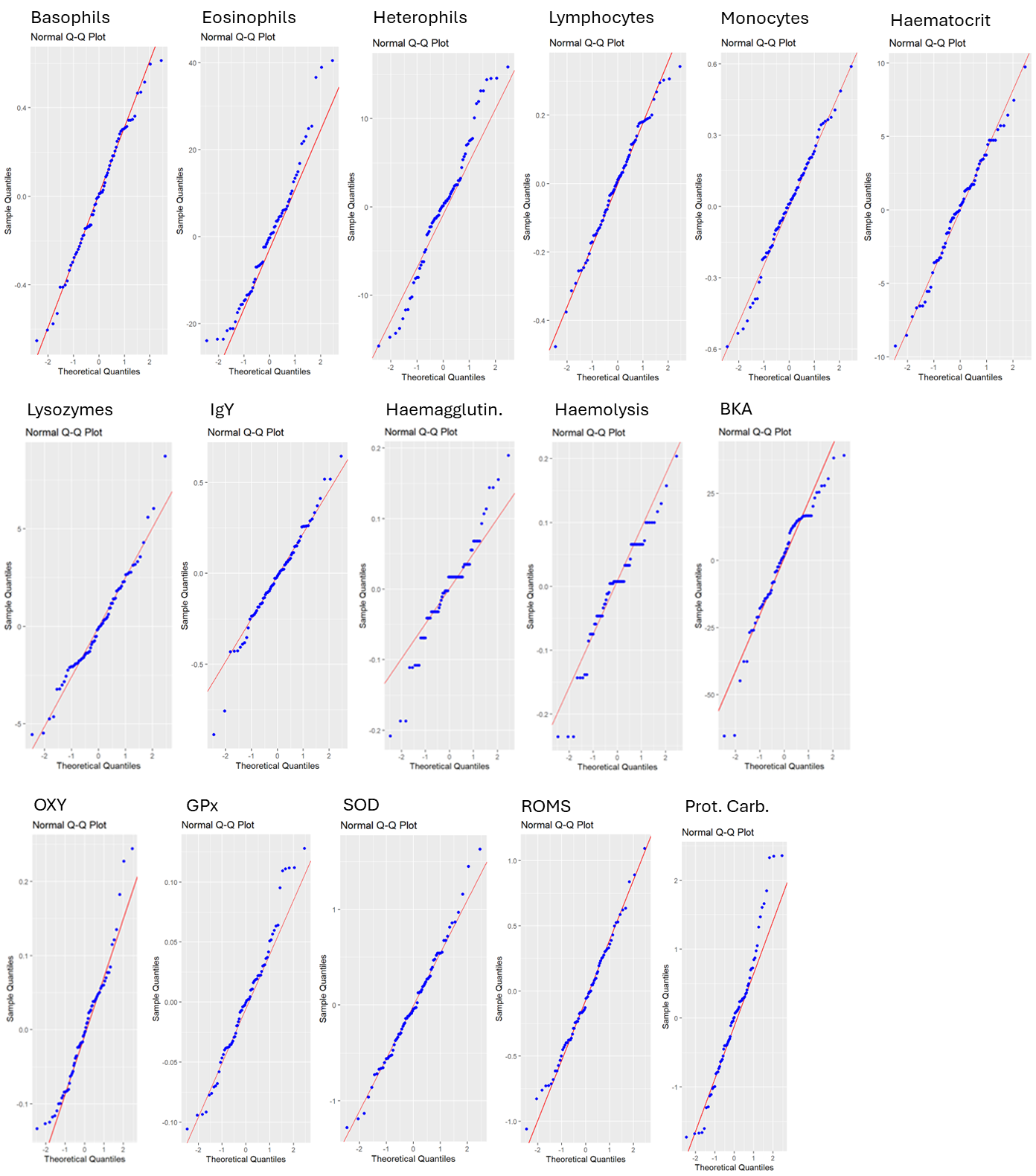

Fig. S2 –Q-Q plots showing model residual distribution for each immune-physiological variable after data transformation and outliers’ removal (best models). Normal distribution of residuals was used to assess the validity of our models.

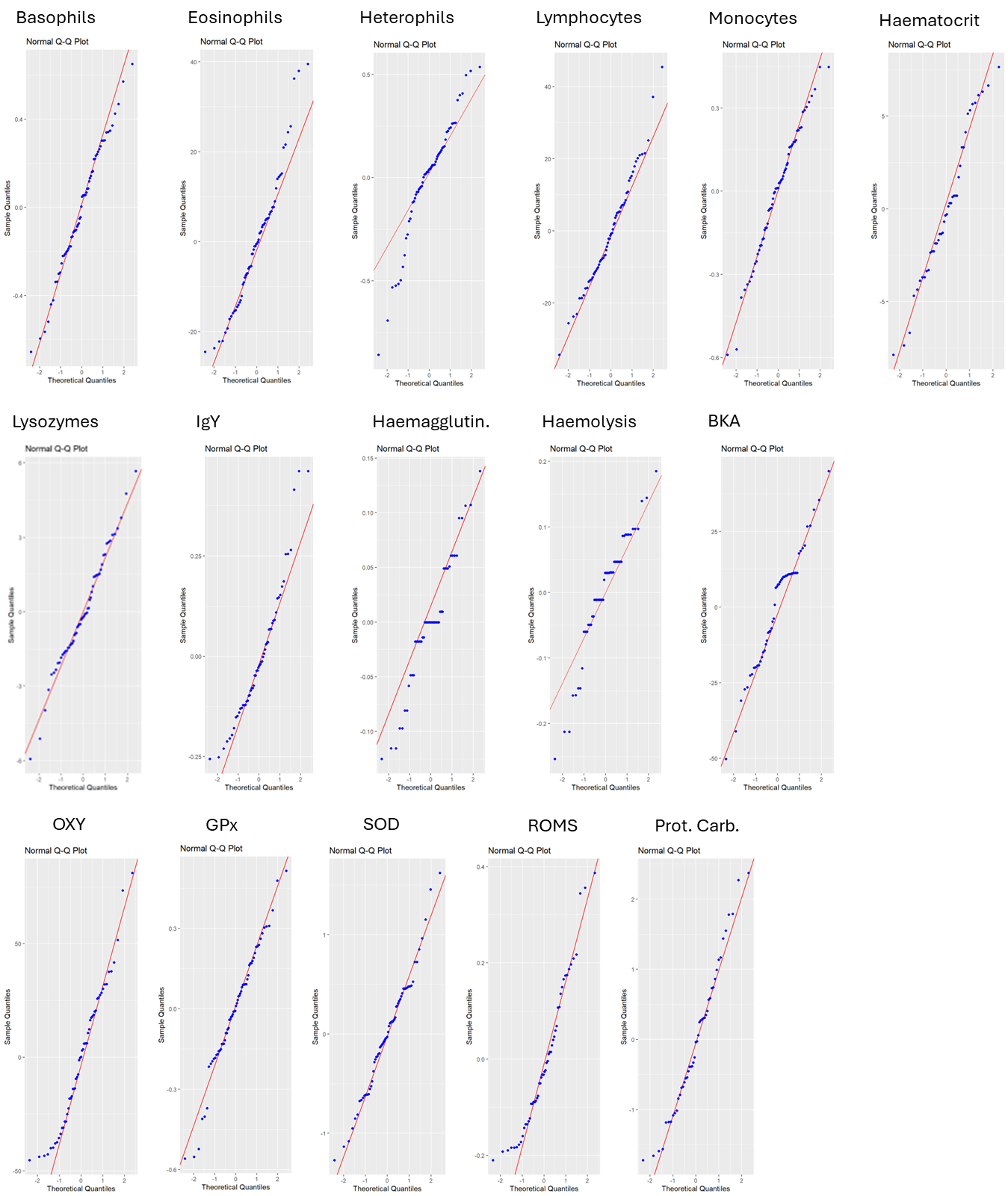

Fig. S3 –Q-Q plots showing residual distribution of models testing the interaction between species and sex. Normal distribution of residuals was used to assess the validity of our models.

Table S1 – Results of LMs including outliers. ALEG = Egyptian goose; ANPL = mallard; CYOL = mute swan. Significant P-values are indicated in bold.

| **Marker** | **Variable tranformation** | **Contrast** | **Estimate** | **SE** | **t value** | **P-value** |
| --- | --- | --- | --- | --- | --- | --- |
| Basophils | Log | ALEG - ANPL | -0.76 | 0.23 | -3.28 | **<0.01** |
|  |  | ALEG - CYOL | -0.32 | 0.16 | -2.02 | 0.05 |
|  |  | CYOL - ANPL | -0.44 | 0.22 | -1.96 | 0.05 |
| Eosinophils | None | ALEG - ANPL | -6.15 | 5.73 | -1.07 | 0.29 |
|  |  | ALEG - CYOL | 1.79 | 3.92 | 0.46 | 0.65 |
|  |  | CYOL - ANPL | -7.93 | 5.56 | -1.43 | 0.16 |
| Heterophils | None | ALEG - ANPL | -4.19 | 3.10 | -1.35 | 0.18 |
|  |  | ALEG - CYOL | -9.71 | 2.12 | -4.59 | **<0.01** |
|  |  | CYOL - ANPL | 5.52 | 3.01 | 1.84 | 0.07 |
| Lymphocytes | Log | ALEG - ANPL | -0.18 | 0.07 | -2.74 | **0.01** |
|  |  | ALEG - CYOL | -0.16 | 0.04 | -3.66 | **<0.01** |
|  |  | CYOL - ANPL | -0.02 | 0.06 | -0.24 | 0.81 |
| Monocytes | Log | ALEG - ANPL | 0.04 | 0.09 | 0.45 | 0.66 |
|  |  | ALEG - CYOL | 0.19 | 0.06 | 2.95 | **<0.01** |
|  |  | CYOL - ANPL | -0.15 | 0.09 | -1.62 | 0.11 |
| Haematocrit | None | ALEG - ANPL | 4.22 | 3.09 | 1.37 | 0.18 |
|  |  | ALEG - CYOL | 4.89 | 2.15 | 2.27 | **0.03** |
|  |  | CYOL - ANPL | -0.67 | 3.14 | -0.21 | 0.83 |
| Lysozymes | None | ALEG - ANPL | -0.82 | 0.97 | -0.85 | 0.40 |
|  |  | ALEG - CYOL | -4.00 | 0.68 | -5.90 | **<0.01** |
|  |  | CYOL - ANPL | 3.17 | 0.94 | 3.36 | **<0.01** |
| Haemagglutination | Log | ALEG - ANPL | -0.03 | 0.04 | -0.92 | 0.36 |
|  |  | ALEG - CYOL | -0.06 | 0.03 | -2.33 | **0.02** |
|  |  | CYOL - ANPL | 0.03 | 0.03 | 0.82 | 0.41 |
| Haemolysis | None | ALEG - ANPL | -0.02 | 0.04 | -0.55 | 0.59 |
|  |  | ALEG - CYOL | -0.08 | 0.03 | -2.70 | **0.01** |
|  |  | CYOL - ANPL | 0.05 | 0.04 | 1.50 | 0.14 |
| IgY | None | ALEG - ANPL | -0.26 | 0.08 | -3.34 | **<0.01** |
|  |  | ALEG - CYOL | -0.31 | 0.05 | -5.64 | **<0.01** |
|  |  | CYOL - ANPL | 0.05 | 0.07 | 0.67 | 0.50 |
| BKA | None | ALEG - ANPL | -0.40 | 8.96 | -0.05 | 0.96 |
|  |  | ALEG - CYOL | 36.00 | 6.59 | 5.46 | **<0.01** |
|  |  | CYOL - ANPL | -36.40 | 8.96 | -4.06 | **<0.01** |
| OXY | None | ALEG - ANPL | 16.86 | 16.80 | 1.00 | 0.32 |
|  |  | ALEG - CYOL | -10.22 | 11.86 | -0.86 | 0.39 |
|  |  | CYOL - ANPL | 27.08 | 16.44 | 1.65 | 0.10 |
| GPx | Log | ALEG - ANPL | 0.09 | 0.09 | 0.99 | 0.33 |
|  |  | ALEG - CYOL | -0.07 | 0.06 | -1.10 | 0.28 |
|  |  | CYOL - ANPL | 0.16 | 0.09 | 1.79 | 0.08 |
| SOD | None | ALEG - ANPL | -0.35 | 0.24 | -1.49 | 0.14 |
|  |  | ALEG - CYOL | -0.06 | 0.16 | -0.36 | 0.72 |
|  |  | CYOL - ANPL | -0.29 | 0.23 | -1.28 | 0.21 |
| ROMS | None | ALEG - ANPL | -0.35 | 0.09 | -4.06 | **<0.01** |
|  |  | ALEG - CYOL | -0.11 | 0.06 | -1.87 | 0.07 |
|  |  | CYOL - ANPL | -0.24 | 0.08 | -2.93 | **<0.01** |
| Protein carbonyls | None | ALEG - ANPL | 0.02 | 0.48 | 0.03 | 0.97 |
|  |  | ALEG - CYOL | 0.55 | 0.34 | 1.64 | 0.11 |
|  |  | CYOL - ANPL | -0.54 | 0.45 | -1.19 | 0.24 |

Table S2 – Results of the ANOVA of LMs testing for the interaction between species and sex. Significant P-values are indicated in bold.

| **Marker** | **Variable tranformation** | **Contrast** | **df** | **Sum sq** | **Mean Sq** | **F-value** | **P-value** |
| --- | --- | --- | --- | --- | --- | --- | --- |
| Basophils | Log | species | 1 | 0.08 | 0.08 | 0.92 | 0.34 |
|  |  | sex | 1 | 0.24 | 0.24 | 2.69 | 0.11 |
|  |  | species*sex | 1 | 0.02 | 0.02 | 0.20 | 0.66 |
| Eosinophils | None | species | 1 | 50.20 | 50.17 | 0.22 | 0.64 |
|  |  | sex | 1 | 38.00 | 37.98 | 0.17 | 0.68 |
|  |  | species*sex | 1 | 11.90 | 11.88 | 0.05 | 0.82 |
| Heterophils | Log | species | 1 | 1.61 | 1.61 | 19.27 | **>0.01** |
|  |  | sex | 1 | 0.36 | 0.36 | 4.35 | **0.04** |
|  |  | species*sex | 1 | 0.18 | 0.18 | 2.19 | 0.14 |
| Lymphocytes | None | species | 1 | 3401.20 | 3401.20 | 14.09 | **>0.01** |
|  |  | sex | 1 | 3857.40 | 3857.40 | 15.98 | **>0.01** |
|  |  | species*sex | 1 | 50.30 | 50.30 | 0.21 | 0.65 |
| Monocytes | Log | species | 1 | 0.67 | 0.67 | 12.11 | **>0.01** |
|  |  | sex | 1 | 0.08 | 0.08 | 1.51 | 0.22 |
|  |  | species*sex | 1 | 0.07 | 0.07 | 1.27 | 0.26 |
| Haematocrit | None | species | 1 | 557.36 | 557.36 | 32.77 | **>0.01** |
|  |  | sex | 1 | 215.46 | 215.46 | 12.67 | **>0.01** |
|  |  | species*sex | 1 | 19.30 | 19.30 | 1.13 | 0.29 |
| Lysozymes | None | species | 1 | 185.48 | 185.48 | 34.22 | **>0.01** |
|  |  | sex | 1 | 3.81 | 3.81 | 0.70 | 0.41 |
|  |  | species*sex | 1 | 27.41 | 27.41 | 5.06 | **0.03** |
| Haemagglutination | Log | species | 1 | 0.08 | 0.08 | 21.58 | **>0.01** |
|  |  | sex | 1 | 0.01 | 0.01 | 2.50 | 0.12 |
|  |  | species*sex | 1 | 0.01 | 0.01 | 1.99 | 0.17 |
| Haemolysis | Log | species | 1 | 0.10 | 0.10 | 10.53 | **>0.01** |
|  |  | sex | 1 | 0.01 | 0.01 | 1.27 | 0.27 |
|  |  | species*sex | 1 | 0.00 | 0.00 | 0.32 | 0.57 |
| IgY | None | species | 1 | 1.52 | 1.52 | 51.34 | **>0.01** |
|  |  | sex | 1 | 0.17 | 0.17 | 5.84 | **0.02** |
|  |  | species*sex | 1 | 0.00 | 0.00 | 0.06 | 0.81 |
| BKA | None | species | 1 | 22077.10 | 22077.10 | 53.64 | **>0.01** |
|  |  | sex | 1 | 360.80 | 360.80 | 0.88 | 0.35 |
|  |  | species*sex | 1 | 367.70 | 367.70 | 0.89 | 0.35 |
| OXY | None | species | 1 | 134.00 | 134.28 | 0.14 | 0.71 |
|  |  | sex | 1 | 2130.00 | 2129.91 | 2.17 | 0.15 |
|  |  | species*sex | 1 | 810.00 | 810.39 | 0.83 | 0.37 |
| GPx | Log | species | 1 | 0.08 | 0.08 | 1.45 | 0.23 |
|  |  | sex | 1 | 0.21 | 0.21 | 3.77 | 0.06 |
|  |  | species*sex | 1 | 0.06 | 0.06 | 1.05 | 0.31 |
| SOD | None | species | 1 | 0.28 | 0.28 | 0.74 | 0.39 |
|  |  | sex | 1 | 0.09 | 0.09 | 0.24 | 0.63 |
|  |  | species*sex | 1 | 0.16 | 0.16 | 0.43 | 0.52 |
| ROMS | None | species | 1 | 0.09 | 0.09 | 3.58 | 0.06 |
|  |  | sex | 1 | 0.00 | 0.00 | 0.07 | 0.80 |
|  |  | species*sex | 1 | 0.02 | 0.02 | 0.93 | 0.34 |
| Protein carbonyls | None | species | 1 | 1.81 | 1.81 | 1.54 | 0.22 |
|  |  | sex | 1 | 0.45 | 0.45 | 0.38 | 0.54 |
|  |  | species*sex | 1 | 0.34 | 0.34 | 0.29 | 0.59 |

Table S3 – Pairwise comparison of the significant interaction term for lysozymes. ALEG = Egyptian goose; CYOL = mute swan; f = female; m = male. Significant P-values are indicated in bold.

| **Marker** | **Variable tranformation** | **Contrast** | **Estimate** | **SE** | **df** | **t. ratio** | **P-value** |
| --- | --- | --- | --- | --- | --- | --- | --- |
| Lysozymes | Log | ALEG f - ALEG m | -0.94 | 0.90 | 55 | -1.04 | 0.73 |
|  |  | CYOL f - CYOL m | 1.87 | 0.87 | 55 | 2.16 | 0.15 |
|  |  | ALEG f - CYOL f | -4.72 | 0.84 | 55 | -5.61 | **<0.01** |
|  |  | ALEG m - CYOL m | -1.91 | 0.92 | 55 | -2.07 | 0.18 |
